# Stability and volatility shape the gut bacteriome and mycobiome dynamics in a pig model

**DOI:** 10.1101/2022.02.02.478893

**Authors:** Brandi Feehan, Qinghong Ran, Victoria Dorman, Kourtney Rumback, Sophia Pogranichniy, Kaitlyn Ward, Robert Goodband, Megan C Niederwerder, Katie Lynn Summers, Sonny T M Lee

## Abstract

The gut microbiome plays important roles in the maintenance of health and pathogenesis of diseases in the growing host. In order to fully comprehend the interplay of the gut microbiome and host, a foundational understanding of longitudinal bacteriome and mycobiome development is necessary. In this study, we evaluated enteric bacterial, fungal and host dynamics throughout the lifetime of commercial swine. We collected a total of 234 fecal samples from 9 pigs across 31 time points in 3 developmental stages (5 preweaning, 15 nursery, and 11 growth adult). We then performed 16S rRNA amplicon sequencing for bacterial profiles and qPCR for the fungus *Kazachstania slooffiae*. We identified distinct bacteriome clustering according to the host developmental stage, with the preweaning stage exhibiting low bacterial diversity and high volatility amongst samples. We further recovered clusters of bacterial populations that were considered core, transient and increasing throughout the host lifetime, suggesting distinct yet important roles by which these differing bacterial population clusters played in the different host stages. *Kazachstania slooffiae* was absent in the preweaning stage but peaked during the nursery stage of the host. We determined that all host growth stages contained negative correlations between *K. slooffiae* and bacterial genera, with only the growth adult stage containing positive correlates. The lack of positive correlates and shared *K. slooffiae-bacteria* interactions between stages warrants future research into the interactions amongst these kingdoms for host health. This research is foundational for understanding how the bacteriome and mycobiome develop singularly, as well as within a complex ecosystem in the host’s gut environment.

## Introduction

Host-associated microbiomes have critical roles in host health, growth and development. The digestive system contains microbes with a wide array of functions for hosts, such as aiding in nutrient availability, protecting from pathogen invasion and maintaining a healthy gut epithelial barrier (Barko et al., 2018; Lloyd-Price et al., 2016; Niederwerder, 2018). An imbalance of microorganisms, or their associated functions, in this enteric, or digestive, microbiome can lead to a dysbiotic state and diseased host (Barko et al., 2018). Diseases and symptoms associated with a dysbiotic enteric microbiome include inflammatory bowel disease (IBD), diarrhea, obesity, and metabolic syndrome (MetS), among other ailments (Pushpanathan et al. 2019). In order to develop therapies for these illnesses, it is paramount to understand the enteric microbiome dynamics spanning microbial kingdoms, including bacteria and fungi, throughout the lifetime of asymptomatic hosts.

Understanding microbial correlations and interactions between fungi and bacteria are critical to elucidating diseases impacted by these microbial kingdoms. Previous research has shown a negative correlation, indicating a competitive relationship, between bacterial diversity and fungal abundance (Kapitan et al. 2019). More specifically, intestinal bacteria have been demonstrated as mediating colonization resistance against *Candida albicans* in mice (Gutierrez et al. 2020). Researchers hypothesize that enteric fungi, especially *Candida* species, appear to be opportunistic as they increase in abundance when bacterial species decline (Kapitan et al. 2019). Still, the microbial mechanisms, influencing other microbes and the host alike, underlying these outcomes have not been described. We must understand bacterial-fungal interaction intricacies in order to provide treatments targeting specific microbes and mechanisms, especially those of bacterial-fungal dysbiotic gut microbiomes.

This study highlights development of the bacteriome, mycobiome and host, with a thorough investigation into correlations between the two microbial kingdoms. This study followed 9 apparently healthy swine from birth to adult. Swine are a suitable model for humans since they have similar diets, genetic content, gastrointestinal systems and immune systems, along with other characteristics (Wang and Donovan 2015). Understanding inter-kingdom interaction in the swine host may thus provide insights into the intricate relationship between the human host and the microbiome. Foundation to this longitudinal study, swine were grown in three stages which varied according to host development, diet and housing: preweaning (milk diet with littermates and dam; birth-21 days of age), nursery (pellet diet co-housed with other litters; 21-80 days) and growth adult (pellet diet co-housed with other litters; 80-122 days). Directly following weaning into the nursery stage in swine hosts, one fungus is consistently identified in the enteric mycobiome: *Kazachstania slooffiae* (Summers et al. 2019; Arfken et al. 2019). For this reason, our study focussed on elucidating longitudinal dynamics between *K. slooffiae* and bacteria.

*Kazachstania slooffiae* is a member of the Saccharomycetaceae family, and the fungus is a proposed commensal in the swine gut microbiome (Summers et al. 2021). Previous studies have identified eight correlations between *K. slooffiae* and microbial genera from nursery-aged hosts (Arfken et al. 2020; Arfken et al. 2019). We hypothesized there were more inter-kingdom correlations occurring throughout the lifetime of the hosts which influence microbiome establishment and host health (Kapitan et al. 2019). Our study aimed to elucidate novel stage-associated bacteriome-mycobiome correlations to build a foundation for future inter-kingdom interaction studies.

In our study, we determined specific host-age and -dietary stage microbiome development characteristics. These included an increasing bacteriome diversity, decreasing bacteriome volatility and increasing mycobiome in the young host (preweaning and nursery developmental stages). The older host (growth adult stage) microbiome was relatively established with a complex correlation network amongst bacteria and *K. slooffiae*. Together, these findings indicated a dynamic microbiome development from birth until weaning with an increasing number of inter-kingdom interactions throughout the host lifetime.

## Materials and Methods

### Hosts and Study Design

We followed 9 swine over the course of their lifetime, from 2 days to 157 days of age, to understand successive shifts in microbial populations in a model system (Supplementary Table S1). The hosts were housed indoors and fed distinct diets according to their stage of life (Supplementary Table S1). Hosts were sampled in three stages: preweaning, nursery, and growth. Preweaning diet consisted of mother’s milk and potentially feed as the hosts grew old enough to reach their mother’s trough. Nursery diet, phase 1, transitioned from milk to pelleted feed after weaning from the mother and moving into a new barn environment. A second pelleted feed was fed during nursery phase 2, while a meal was fed for nursery phase 3. The growth adult stage also included three phase diets with an initial move into another barn environment accompanying the nursery-growth transition (Supplementary Table S1). Hosts did not receive antibiotics or antifungals prior to or during the study.

We used a fresh set of sterile gloves to collect each fecal sample free-catch, prior to contact with the ground. We collected fecal samples every five days during preweaning and nursery stages, and every seven days during the growth adult stage. Immediately after collection, samples were stored in either a sterile 15ml tube or sterile bag, kept on ice, and then transported to the laboratory for subsequent storage at −80°C until genomic DNA extraction.

### DNA Extraction and Marker Gene Sequencing

We used the E.Z.N.A.^®^ Stool DNA Kit (Omega Bio-tek, Inc.; Norcross, GA) to extract the microbial DNA from the fecal samples according to the manufacturer protocols. Extracted DNA was quantified with Nanodrop and a Qubit™ dsDNA BR Assay Kit for sample DNA quality and concentration. Extracted microbial DNA was stored at −80°C until library preparation. Bacterial 16S rRNA V4 region was amplified during library preparation via Illumina’s Nextera XT Index Kit v2 (Illumina, Inc.; San Diego, CA) (primers: 515F, GTGCCAGCMGCCGCGGTAA and 806R, GGACTACHVGGGTWTCTAAT) (Caporaso et al., 2011). Sequencing was done on the Illumina MiSeq which generated paired-end 250bp reads.

### Kazachstania slooffiae qPCR

We performed the *K. slooffiae* qPCR, utilizing the SensiMix™ SYBR^®^ Hi-ROX Kit (Bioline, Meridian Bioscience; Cincinnati, OH), as previously described (primers: KS-f, ATCCGGAGGAATGTGGCTTC and KS-r, AGCATCCTTGACTTGCGTCG) (Urubschurov et al. 2015). Master mix components and qPCR conditions are listed in Supplementary Table S2. Each qPCR run included at least one PCR-grade water with the master mix as a non-template control (NTC), with *K. slooffiae* sample as the positive control.

### Bioinformatic and Statistical Analysis

We used cutadapt and DADA2 in QIIME2 v2019.7 (Callahan et al. 2016) to trim and perform quality control for the sequencing reads (Supplementary Table S3). Reads in which no primer was found were discarded. The reads were truncated at locations where 25 percentile of the reads had a quality score below 15. Diversity analysis was carried out at a sampling depth of 11,105 reads. The pre-trained classifier offered by QIIME2 using the Silva 132 database was used for taxonomic assignment for bacteria. We used a weighted UniFrac on the rarefield dataset (11,105 reads) to evaluate differential microbial composition among the samples in different stages, and used a principal coordinate analysis (PCA) to visualize the microbial composition structure. We calculated the α-diversity to represent the species diversity in each sample, and used Shannon and Faith’s phylogenetic diversity indices to measure the number of species and the uniformity of species abundance to evaluate the species diversity.

We used PERMANOVA in the R Adonis package to test if there were any statistical differences among the stages, and which bacterial populations were distinct to their respective growth stages. We further used DESeq2 to mark the statistical differences in the bacterial populations (phyla and genera) predominance among the stages. 16S rRNA amplicon genera and fungal qPCR Ct values were utilized in a SPIEC-EASI co-occurrence network analysis as previously performed (Arfken et al. 2020). Specific SPIEC-EASI parameters included: Meinhausen–Buhlmann estimation method, lambda minimum ratio of 0.01, and nlambda of 20 (Kurtz et al. 2015). All raw sequence data were deposited in NCBI under the project PRJNA798835.

## Results and Discussion

We collected a total of 234 samples across 31 time points (5 preweaning, 15 nursery, and 11 growth) from 9 pigs (Supplementary Table S1). A total of 10,187,636 sequences resulted from sequencing; we recovered an average of 33,394 ASVs per sample following QIIME2 quality control. Out of the recovered ASVs, an average of 80.1% (79.7% bacteria and 0.4% archaea) populations were annotated on SILVA (Supplementary Table S4).

### Volatility in the preweaning stage led to microbial establishment and stability in later growth stages

As shown in Figure 1A, the weighted UniFrac PCA illustrated a distinct clustering of bacterial community composition between the three growth stages as the pig transitioned from young host to adult. We further observed that there were three distinct clusters of the swine hosts in the nursery stage due to their diet, with the latter diet-stage of nursery being more similar to the growth adult hosts. We showed in our study that a young host lacked an established, shared microbiome, but converged with age and environmental changes such as diet and shared housing. The preweaning host had the most diverse microbial composition amongst the individuals, while the nursery aged host was primarily clustered with the growth host suggesting a rather stable microbiome once the microbial members had established (Slifierz et al., 2015; Wang et al., 2019). Previous research has illustrated distinct microbial populations in swine hosts according to stage of development (Kim et al., 2015; Slifierz et al., 2015; Wang et al., 2019).

**Figure 1:**
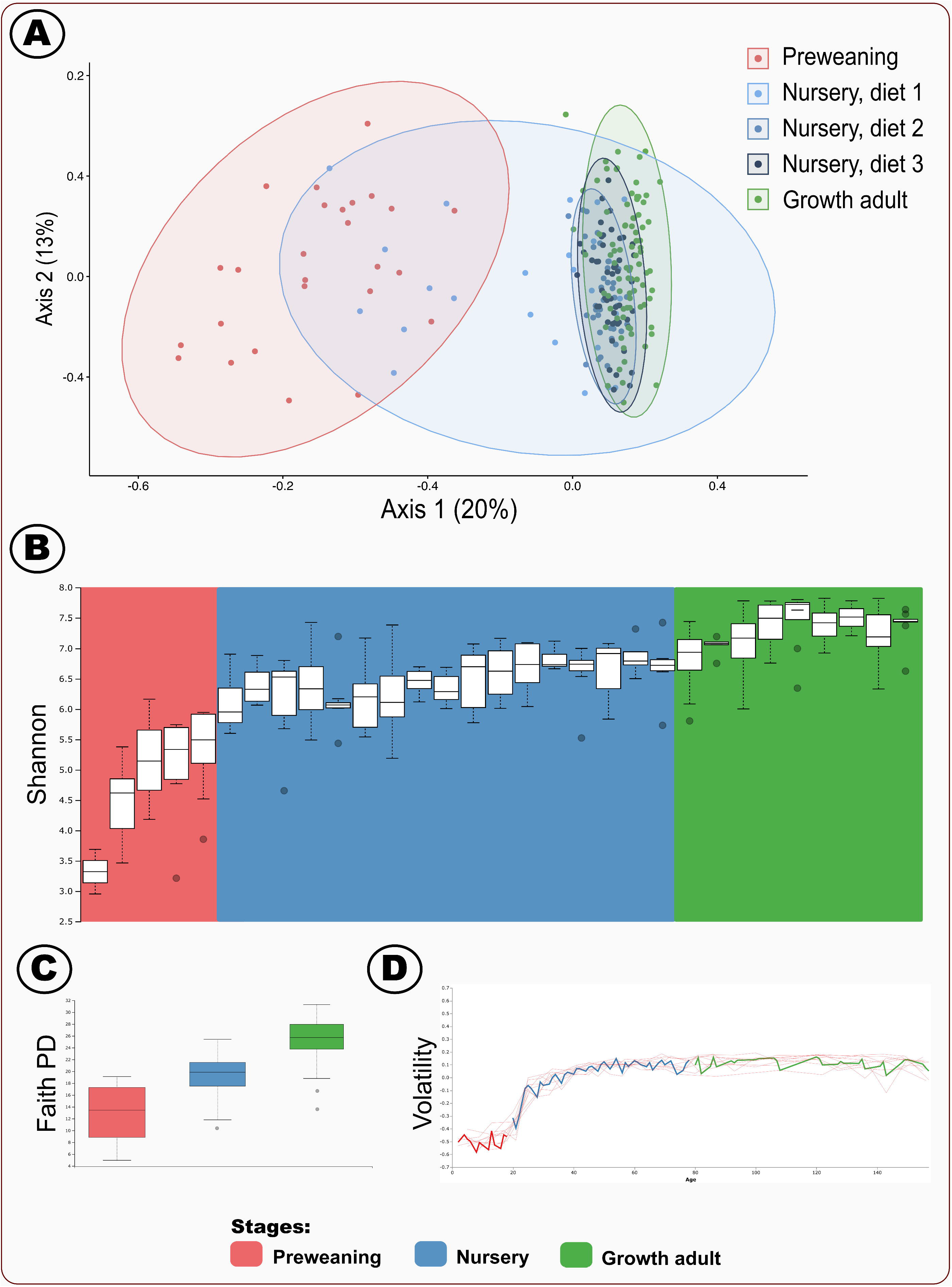
A) Weighted uniFrac PCA plot depicting -diversity; dots represent distinct samples. Nursery stage is separated according to the three diets fed during the stage. B) Longitudinal Shannon diversity. C) Faith’s phylogenetic diversity (PD). D) Volatility control chart of the first axis of the PCA (A).

Our data showed that the Shannon diversity (*p* = 7.7 x 10^−18^) and Faith’s phylogenetic diversity (preweaning (P) vs nursery (N): *p* = 2.5×10^−11^; N vs growth adult (G): *p* = 1.2×10^−18^; P vs G: *p* = 7.4×10^−14^) indices paralleled the PCA analysis (Figure 1B and Figure 1C). We observed that the nursery and growth hosts showed relatively similar levels of microbial diversity, whereas the preweaning host demonstrated comparatively lower diversity which increased until weaning. A second increase, albeit smaller, in Shannon diversity occurred in the growth adult hosts. Faith’s phylogenetic diversity further corroborated the increasing diversity from the preweaning host to growth host. Studies have indicated the increase in microbial diversity during host development is typical across many different host species (Barko et al., 2018; Lloyd-Price et al., 2016).

The microbial composition volatility index in the preweaning host hovered near −0.5 while approaching 0 in the early nursery stage (Figure 1D). These volatility findings further suggested that the young preweaning host had a relatively more volatile, fluctuating microbiome. Our results were consistent with other mammalian studies that demonstrated a volatile youth microbiome establishment period in children aged from birth to approximately 3 years of age (Yatsunenko et al. 2012).

Putting together the PCA, diversity indices and volatility analyses, our study suggested that the preweaning neonate host had a developing gut microbiome. We showed that the microbiome was converging in the late preweaning and early nursery host, and there were relatively little changes occurring in microbial diversity after the convergence of the microbial community in the intermediate nursery host.

### Foundational and transient bacterial populations in different growth stages

We analyzed the host microbial membership and identified 23 distinct phyla (Figure 2, Supplementary Table S5). We demonstrated a core bacterial population consisting of seven phyla which dominated throughout the lifetime of the swine host, suggesting these bacterial populations have essential implications to the host’s health and well-being (Ke et al., 2019; Kumar et al., 2019/5; X. Wang et al., 2019). Our study showed that Bacteroidetes and Firmicutes were the predmoninating core microbes throughout the host lifetime (Figure 2A). These results were consistent with findings from previous research demonstrating similar development of increasing bacterial diversity, with Firmicutes and Bacteroidetes consistently dominating (Ke et al., 2019; Kumar et al., 2019/5; Wang et al., 2019). We also noticed that clustered alongside these two core bacterial phyla, albeit at relatively lower abundances, included Spirochaetes, Epsilonbacteraeota, Proteobacteria, Tenericutes, and Actinobacteria (Figure 2A). With Epsilonbacteraeota being an exception, the above seven phyla have been identified consistently in human and swine hosts throughout their lifetimes (T. Chen et al., 2019; Niu et al., 2015). Although Spirochaetes (Nakamura, 2020; Neef et al., 1994), Epsilonbacteraeota (He et al., 2020) and Proteobacteria (Ghanbari et al., 2019; Torres Luque et al., 2020) have been classified as pathogenic opportunistic organisms in previous studies, more research is necessary to understand their role in modulating gut health and dysbiosis (He et al., 2020; Nakamura, 2020; Neef et al., 1994).

**Figure 2:**
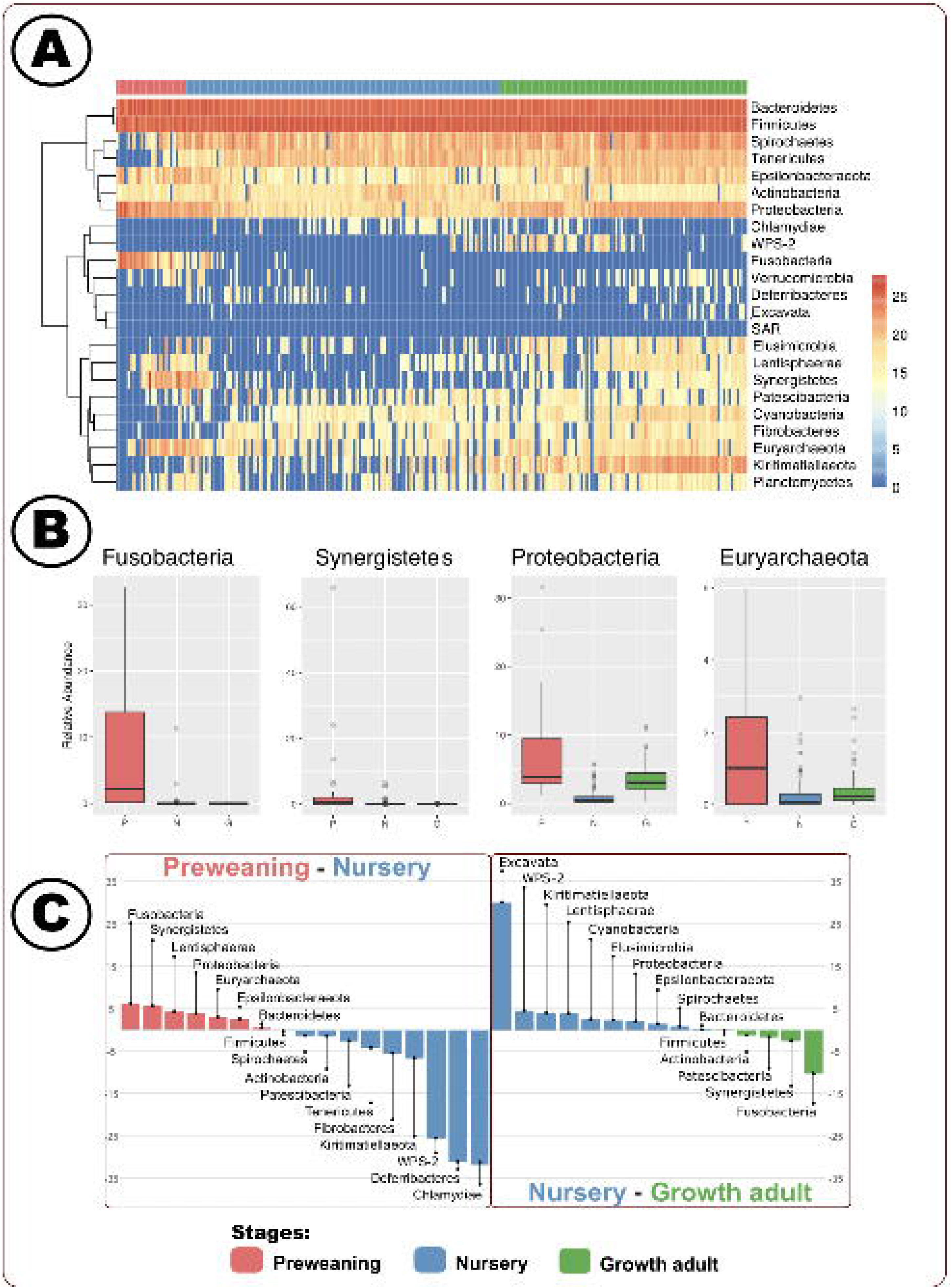
A) Longitudinal heat map of DESeq2 resulting phyla relative abundances. B) Relative abundance of select phyla. C) DESeq2 differentially identified (p<0.05) phyla.

Beyond the seven core phyla, we also identified a set of bacterial populations that demonstrated distinct development patterns - bacterial phyla identified only in the preweaning and growth adult hosts, and phyla which were increasing in relative abundance alongside host age (Figure 2A). We surmised that these bacterial populations were transitive at different stages of the host. We noticed that Elusimicrobia, Lentisphaerae, and Synergistes only existed in the preweaning and growth adult, but not in the nursery stages of the host. On the other hand, phyla that were increasing in relative abundance throughout the host lifetime were Patescibacteria, Cyanobacteria, Fibrobacteres, Euryarchaeota, Kiritimatiellaeota, and Planctomycetes. We suggest that the establishment of microbial populations during the preweaning stage was crucial to the well-being of the host due to the relatively unstable, forming microbiome. Fusobacteria, Synergistetes, and Proteobacteria had higher relative abundance in the young preweaning host as identified 28 significant DESeq2 differential gecompared to the weaned, nursery host (Figure 2B). Fusobacteria have been identified in hosts with infections, obesity, and heat stress (He et al., 2020; Panasevich et al., 2018; Y. Wang et al., 2019). On the other hand, previous studies have found that Synergistetes (Li et al., 2014; Magalhaes et al., 2007) and Proteobacteria (Ghanbari et al., 2019; Torres Luque et al., 2020) can be commensal or pathogenic. Altogether, these results suggested the young host’s gut microbiome was relatively unstable as would be found in a dysbiotic state. Further research is crucial to understand the enteric microbial functional dynamics associated with host dynamics in growth and health.

We noticed that the differences between the nursery and growth adult host were relatively fewer than the preweaning to the nursery host, and the shaping of the microbiome as the host matures might result in commensal and opportunistic pathogens that were stage-dependent. We noticed that Chlamydiae, Deferribacteres and WPS-2 were prominent in the nursery host, suggesting that bacterial populations from these three phyla might play an important role in the host well-being as the swine gut microbiome started to stabilize throughout the stages (Figure 2C). The roles Chlamydiae (Karasova et al., 2021; Szeredi et al., 1996) and WPS-2 (Ward et al., 2019) play in the gut are unclear, as these microbes could be commensal or opportunistic. Likewise, the impact of Deferribacteres are not certain, but Deferribacteres is known to appear according to changes in dietary components including iron (Hu et al., 2020; Ma et al., 2019; Ogita et al., 2020). Although these three phyla were more than 25 times higher in relative abundance in the nursery hosts relative to preweaned hosts, the average relative abundance per phyla per sample in nursery hosts was less than 1% (Figure 2C). The comparatively low relative abundance supported the previous finding of increased diversity in the intermediate aged hosts. While there were potential pathogenic organisms identified in the nursery-aged host, these bacterial populations were found at a relatively low abundance in an increasingly diverse and established gut environment post-weaning.

We also observed that an archaeal phylum, Euryarchaeota, was associated with the preweaned stage. Microbial interactions between archaea and bacteria are paramount to elucidating microbiome development with consequences on host health. We hypothesize that Euryarchaeota might be working alongside and with the bacteria community to shape the host microbiome, which might influence overall host health and well-being (Hillman et al., 2017). Euryarchaeota has been associated with improved fiber digestion (T. Chen et al., 2019; Tang et al., 2020), and have been identified in preweaned swine previously, although the interaction of archaea with bacteria and fungi in swine is not fully understood (Federici et al. 2015).

We identified 28 significant DESeq2 differential genera (*p* < 0.01) amongst the three stages (Figure 3 and Supplementary Table S5). Unlike the phyla level analysis, we did not observe a core genus but instead identified three distinct clusters throughout the lifetime of the host (Figure 3A). The first cluster consisted solely of *Bacteroides* as the bacterial population decreased post-weaning. *Succinivibrio* and *Selenomonas* appeared most abundant at different times in the nursery host followed by a rebound period, consisting of a decrease and final increase to a plateau in the growth adult stage. The final cluster, with 26 genera, followed a similar pattern, but with two prominent peaks (near weaning, intermediate nursery) and a final rebound similar to the second cluster (with the rebound plateau in intermediate growth). Interestingly, *Bacteroides*, *Succinivibrio*, and *Selenomonas* are all heavily reliant on carbohydrate utilization (Flint, 2004; Graf et al., 2015; Sawanon et al., 2011). Previous research has indicated these genera utilize distinct carbohydrate resources (Hailemariam et al., 2020; Hooper et al., 2002; Kenny et al., 2011), but future research is necessary to confirm how the bacterial species in these hosts were utilizing and interacting among the microbes and host, in their use of carbohydrates.

**Figure 3:**
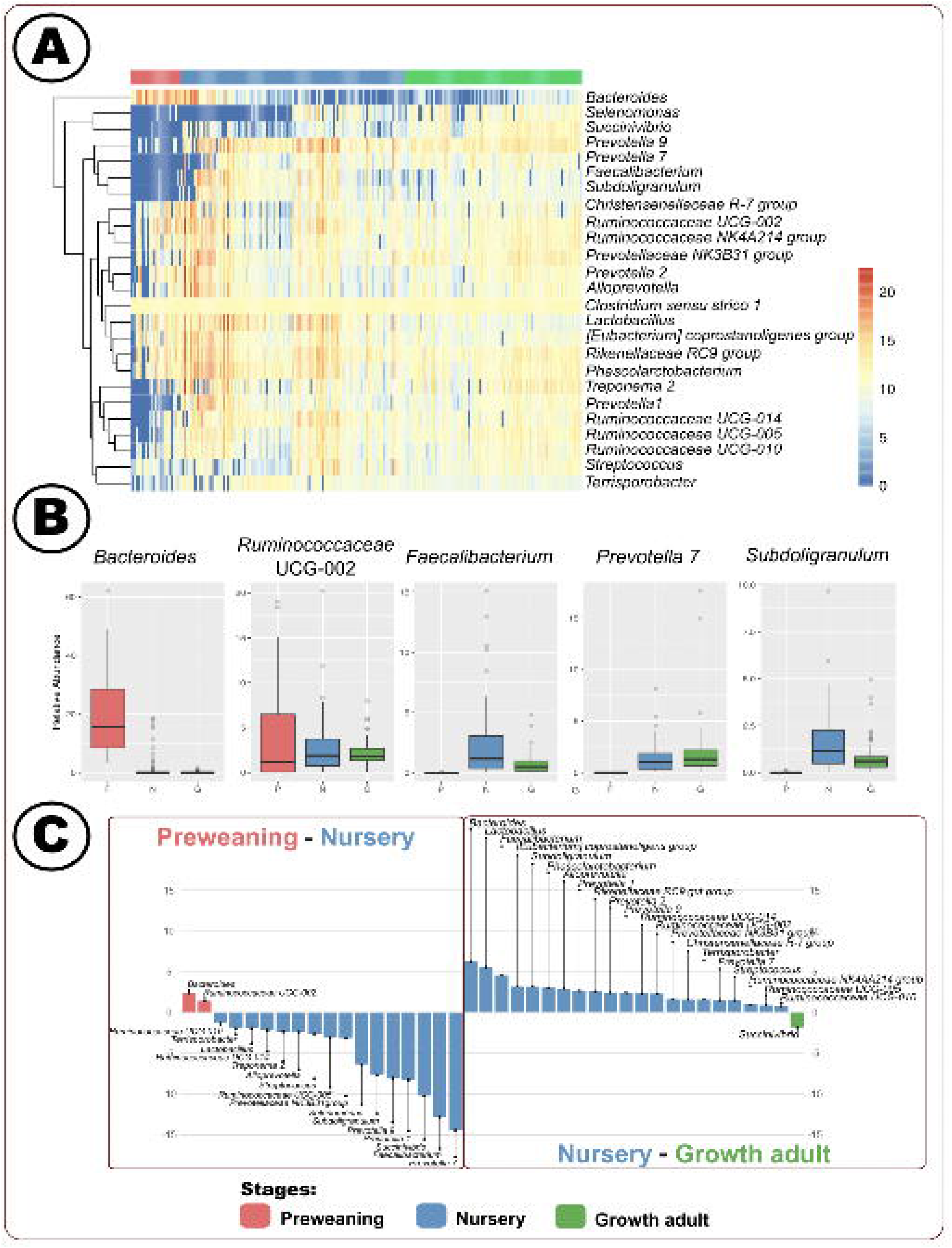
A) Longitudinal heat map of DESeq2 resulting genera relative abundances. B) Relative abundance of select genera. C) DESeq2 differentially identified (p<0.05) genera.

Differential stage analysis indicated *Bacteroides* and *Ruminococcaceae UCG-002* were identified only in preweaning hosts but absent in the intermediate aged hosts (Figure 3B). On the other hand, we determined that *Faecalibacterium, Prevotella 7*, and *Subdoligranulum* had higher relative abundance in the intermediate host as compared to the neonate, suggesting that the microbial function of fermenting dietary carbohydrates into short chain fatty acids (SCFAs) (Arfken et al., 2019; Biddle et al., 2013; De Filippo et al., 2010; Michalak et al., 2020) in these differential genera played an important role during the growth of the host in the nursery and growth adult stages (Figure 3B). Our study further supported previous bacteriome establishment dynamics while elucidating novel stage-associations highlighting a need for functional determination of the enteric microbiome according to host development.

### Temporal dynamics of Kazachstania slooffiae and association with bacterial diversity

Our study suggested that fungal-bacterial interactions in the swine host could influence both bacteriome and mycobime establishment and dynamics, therefore leading to the decline in *K. slooffiae* abundance in hosts. We performed qPCR and demonstrated varied *K. slooffiae* abundance according to developmental stage (Figure 4). We noticed the fungus was absent in the preweaning host but its presence peaked in the nursery host from 25-46 days of age, with a steady decrease in abundance past 46 days of age. We determined fungal presence was more dispersed in the older host, as indicated by a larger 95% confidence interval. Interestingly, we found the increase in *K. slooffiae* coincided with the establishment of the microbiome near weaning. Previous studies have indicated an increase of *K. slooffiae* in the early nursery stage (swine hosts aged 21-35 days) (Arfken et al. 2020; Summers et al. 2019). *K. slooffiae* abundance past 35 days of age were previously unknown. Our findings showed that *K. slooffiae* abundance declined during the late nursery stage and plateauing in the growth adult stage, added to the growing knowledge in the understanding of this fungi. Our fungal research suggested that *K. slooffiae* underwent stage-specific growth patterns, similar to that of the bacteriome. The factors which directly influenced *K. slooffiae* increase and decline are not yet known. Prior publications indicate associations between members within the microbiome, including between fungi and bacteria, and may have implications to the well-being of the hosts (Kapitan et al. 2019).

**Figure 4:**
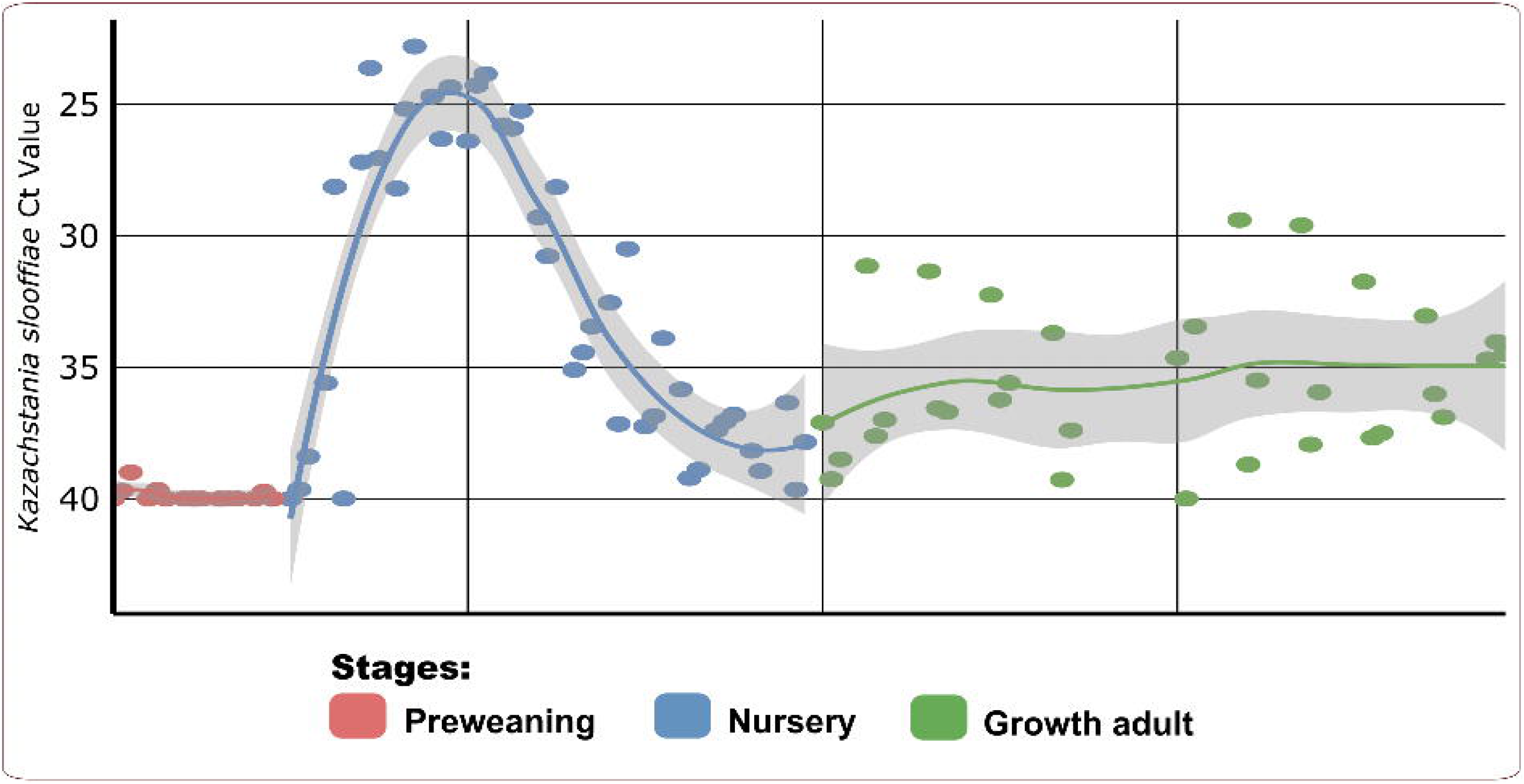
*Kazachstania slooffiae* qPCR Ct value according to day of age with line of best fit and 95% confidence interval by stage.

We performed taxonomic correlation analyses to further investigate fungi-bacteria interactions in the gut microbiome. Our increasing correlation network complexity with host age and lack of shared *K. slooffiae*-correlating genera across stages highlighted stage-dependent microbiome development. We simplified our correlation models to depict direct correlations between *K. slooffiae* and genera according to developmental stage (Figure 5 and Supplementary Table S6). We identified 65 correlations (3 in preweaning, 30 nursery, and 32 growth adult). Previous research has indicated increasingly complex fungal-bacteria network correlations as both the microbiome and host develop from preweaning to nursery, but growth adult stage correlates were previously unknown (Arfken et al. 2020). We identified only two shared between the nursery and growth adult stages: *Rikenellaceae RC9 gut group* and *Candidatus Gastranaerophilales bacterium Zag*. The significance of these genera, especially pertaining to *K. slooffiae*, are not understood and are a topic for future research. The lack of shared *K. slooffiae* correlating taxa may be related to stage-specific bacteria and stage-specific bacteriome-mycobiome interactions.

**Figure 5:**
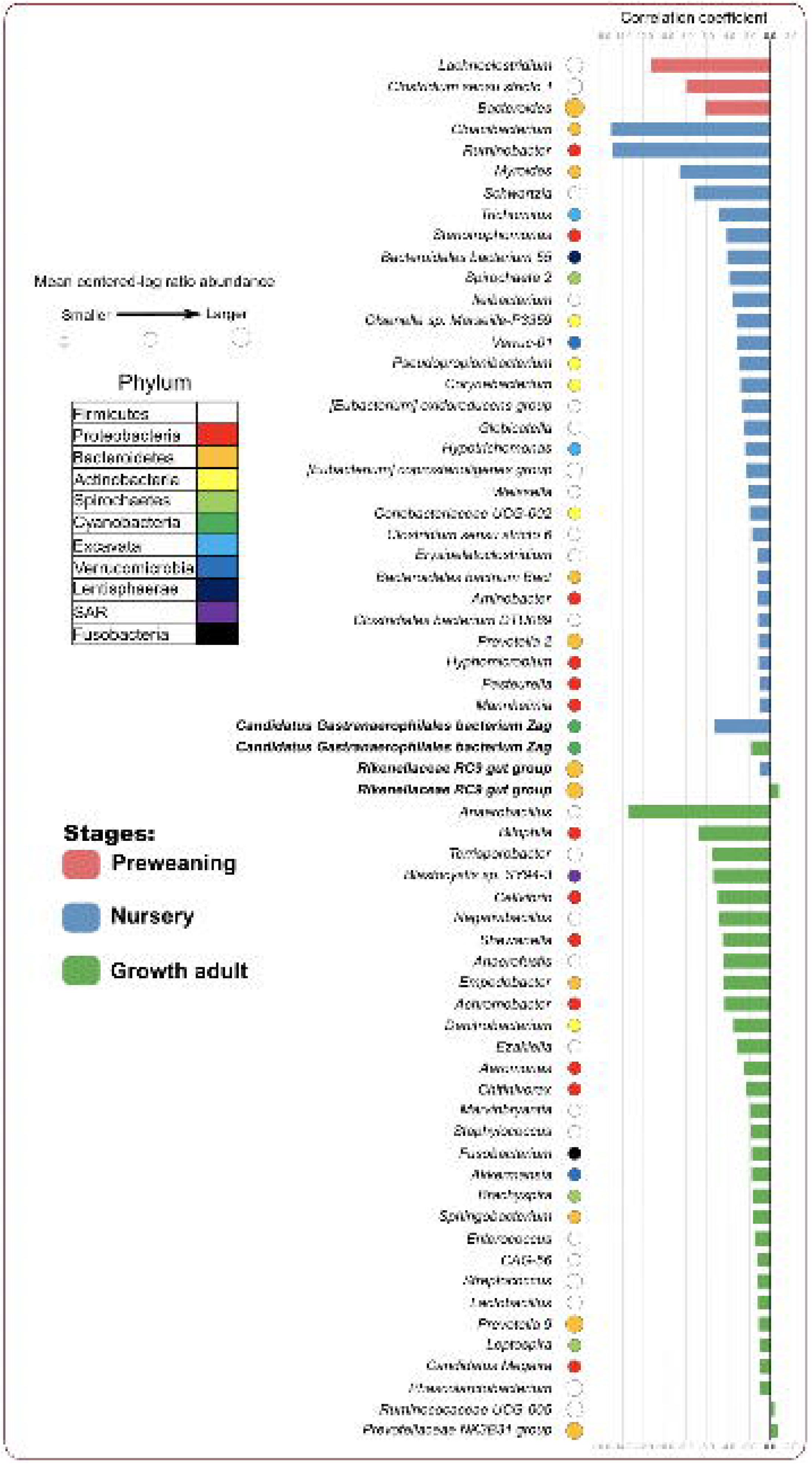
SPIEC-EASI correlation results between *Kazachstania slooffiae* and genera.

Our specific network correlation highlighted novel associations between *K. slooffiae* and the bacteriome throughout the host lifetime, suggesting the changes associated with weaning, including dietary change and stress, may have allowed for *K. slooffiae* expansion while the fungal decline may be attributed to competition with bacteria. Previous publications have identified eight correlations with *K. slooffiae* (Arfken et al. 2020; Arfken et al. 2019). Our results included three out of the eight prior *K. slooffiae* correlations: *Lactobacillus* (correlation coefficient −1.2, growth adult), *Prevotella 9* (−1.2, growth adult), and *Prevotella 2* (−1.1, nursery) (Arfken et al. 2019). Previous research indicated positive correlations of *K. slooffia* and *Lactobacillus, Prevotella 9*, and *Prevotella 2*, whereas our correlations were negative (Arfken et al. 2019). We surmised that negative correlation between *Lactobacillus* and *K. slooffiae* would be analogous to the inhibition of *Lactobacillus* growth by *Candida* in humans (Kapitan et al. 2019). Previous research has identified genetic similarity between *K. slooffiae* and *Candida* (Kurtzman et al. 2005; Summers et al. 2021). Previous studies suggested that *Lactobacillus* may work alongside other bacteria to deter *Candida* growth, such as through short chain fatty acid production (Kapitan et al. 2019). In fact, for our findings, the majority of our network correlations between *K. slooffiae* and genera were negative, with only three positive correlations *(Rikenellaceae RC9 gut group* (0.9), *Prevotellaceae NK3B31 group* (0.7), and *Ruminococcaceae UCG-005* (0.4)) identified in growth hosts. Inverse abundances between fungi and bacteria are indicative of competition or amensalism (Faust and Raes 2012), which could explain the sharp decline of *K. slooffiae* populations in the nursery host (Figure 4). We further hypothesized that the post-weaning increase of *K. slooffiae* might be attributed to the dietary change as *K. slooffiae* is unable to utilize milk galactose (Summers et al. 2021). The dietary transition and host stress from preweaning to nursery might have allowed the increase in *K. slooffiae* populations, even with bacterial establishment relatively progressed (Arfken et al. 2019; Summers et al. 2021). Our correlation network results showed numerous (63 correlations, Figure 5) novel *K. slooffiae* correlations which could aid in divulging establishment dynamics within the bacteriome and mycobiome.

## Conclusions

We provided a comprehensive evaluation of how the bacteriome and the fungus, *Kazachstania slooffiae*, developed through the entire lifetime of a model host, swine. The young preweaning host demonstrated comparatively low microbial diversity which increased near weaning. The growth adult host had a relatively similar microbiome overall compared to the nursery host, yet stage-specific associations, such as potential pathogens and fungal development, were noticed. Our findings provided a foundation for future bacteria-fungi interaction studies.

While microbial inter-kingdom interactions are known to have implications on host health, the intricacies of dynamics between the microbiome and mycobiome are not well understood. Throughout the host lifetime, distinct microbial taxa, diversity, and bacterial-fungi correlations were associated with different stages of life. These stage-associated attributes indicated there could be further stage-associated characteristics such as illness-inducing pathogens and energy providing carbohydrate metabolizing microbes. Future research is crucial to understand the interplay amongst microbes, especially on the functional level pertaining to carbohydrate utilization, according to stage and relating these findings back to host health. Additional research is also necessary to attribute host growth and development, and environmental factors, such as diet and housing, attributing to the diversity changes we identified.

## Acknowledgements

Our team is very grateful to the large number of individuals and organizations which assisted us in performing this research. We thank members of the Kansas State University swine team (Frank Martin, Mark Nelson, Duane Baughman, and Julia Holen) for aiding in the sample collection. Gratitude is also extended to Dr. Alina Akhunova and the Kansas State University Plant Pathology Integrated Genomics Facility for their expertise and assistance in sequencing. Our research was substantially aided by Dr. Ann M. Arfken who assisted in our SPIEC-EASI bacterial-*K. slooffiae* correlation. We greatly appreciate assistance from the following sources: Kansas State University Interdepartmental Genetics Program (fellowship for Brandi Feehan), Global Food Systems Seed Grant Program, Kansas Intellectual and Developmental Disabilities Research Center (NIH U54 HD 090216), the Molecular Regulation of Cell Development and Differentiation – COBRE (P30 GM122731-03) - the NIH S10 High-End Instrumentation Grant (NIH S10OD021743) and the Frontiers CTSA grant (UL1TR002366) at the University of Kansas Medical Center, Kansas City, KS 66160.

## Supplementary

Supplementary Table S1: Demographic and diet composition of the 10 swine hosts.

Supplementary Table S2: *Kazachstania slooffiae* qPCR results, components, and conditions (Urubschurov et al. 2015).

Supplementary Table S3: Raw sequence analysis by QIIME2 Version 2019.7. The table reports on the number of 16S rRNA counts initially obtained, after primer trimming, and DADA2 quality control per sample.

Supplementary Table S4: 16S rRNA amplicon QIIME2 identification counts.

Supplementary Table S5: DESeq2 phyla and genera results comparing preweaning (P), nursery (N) and growth adult (G) stages.

Supplementary Table S6: Raw SPIEC-EASI correlation results.

